# Breast cancer-secreted factors induce atrophy and reduce basal and insulin-stimulated glucose uptake by inhibiting Rac1 activation in rat myotubes

**DOI:** 10.1101/2020.01.27.921262

**Authors:** Mona Sadek Ali, Stine Bitsch-Olsen, Emma Frank, Edmund Battey, Mirela Perla, Scott Sebastian Birch Themsen, Steffen Henning Raun, Steven de Jong, Lykke Sylow

**Affiliations:** Molecular Metabolism in Cancer and Ageing Group, Department of Biomedical Sciences, University of Copenhagen, Denmark; Department of Medical Oncology, Cancer Research Center Groningen, University Medical Center Groningen, University of Groningen, Groningen, The Netherlands

**Keywords:** Breast cancer, Skeletal muscle, Glucose metabolism, Insulin signalling, Muscle atrophy

## Abstract

**Background:** Metabolic disorders are prevalent in women with breast cancer, increasing mortality and cancer recurrence rates. Despite clinical implications, the cause of breast cancer-associated metabolic dysfunction remains poorly understood. Skeletal muscle is crucial for insulin-stimulated glucose uptake, thus key to whole-body glucose homeostasis. In this study, we determined the effect of breast cancer cell-conditioned media on skeletal muscle glucose uptake in response to insulin.

**Method:** L6 myotubes overexpressing myc-tagged GLUT4 (GLUT4myc-L6) were incubated with 40% conditioned media from tumorigenic MCF7 or BT474, or non-tumorigenic control MCF10A breast cells. Mass-spectrometry-based proteomics was applied to detect molecular rewiring in response to breast cancer in the muscle. Expression of myogenesis and inflammation markers, GLUT4 translocation, [^3^H]-2-deoxyglucose (2DG) uptake, and intramyocellular insulin signalling were determined.

**Results:** Breast cancer cell-conditioned media induced proteomic changes in pathways related to sarcomere organisation, actin filament binding, and vesicle trafficking, disrupted myogenic differentiation, activated an inflammatory response via NF-κB, and induced muscle atrophy. Basal and insulin-stimulated GLUT4 translocation and 2DG uptake were reduced in myotubes treated with breast cancer cell-conditioned media compared to the control. Insulin signalling via the Rho GTPase Rac1 was blocked in breast cancer-treated myotubes, while Akt-TBC1D4 signalling was unaffected.

**Conclusion:** Conditioned media from MCF7 and BT474 breast cancer cells reduced skeletal muscle glucose uptake via inhibition of GLUT4 translocation and intramyocellular insulin signalling by selectively blocking Rac1 activation and inducing inflammation. These findings indicate that the rewiring of skeletal muscle proteome, inflammation, and insulin signalling could play a role in metabolic dysfunction in patients with breast cancer.

## Introduction

Breast cancer is the most common type of cancer in the world, responsible for 670.000 deaths in 2022^1–4^. Fortunately, initiatives promoting early cancer detection and improved treatment have resulted in a growing number of breast cancer survivors. However, patients with breast cancer and survivors often display metabolic disorders, including obesity, hyperglycemia, hyperinsulinemia, cardiovascular problems, and glucose intolerance ^5–8^. Consequentially, patients with breast cancer have an increased risk of developing type 2 diabetes after their diagnosis compared to matched individuals without breast cancer^9,10^. Despite great efforts, there is still a lack of knowledge about the physiological and molecular causes of breast cancer-associated metabolic dysfunctions ^5–8,11^. These metabolic dysfunctions increase the risk of mortality^12^ and the likelihood of breast cancer recurrence^13–15^. Thus, gaining insight into these mechanisms could enhance future cancer treatments, ultimately improving long-term survival and overall health.

Metabolic diseases such as type 2 diabetes are largely driven by insulin resistance in skeletal muscle^16,17^. Hence, insulin resistance in skeletal muscle could potentially be the underlying cause of breast cancer-associated metabolic dysfunctions. Skeletal muscle is essential for whole-body metabolic regulation as it is responsible for up to 80% of the glucose uptake in response to insulin^16,17^. Central to insulin resistance is a reduction in insulin-dependent glucose uptake into skeletal muscle, partially due to a reduction in GLUT4 translocation in both rodent^18,19^ and human^20,21^ skeletal muscle. We and others have identified the Rho GTPase Rac1 as a key regulator of skeletal muscle insulin sensitivity^22–27^. This pathway acts in parallel to and mostly independently of insulin-stimulated Akt activation ^24–26,28^. Studies have shown that Rac1 is dysregulated in skeletal muscle of insulin-resistant obese and type 2 diabetic rodents and humans^25–27^.

Breast cancer has been reported to cause metabolic dysfunctions, decrease muscle strength and increase inflammation in skeletal muscle ^29–32^. Recent advances in omics technologies, particularly proteomics, have helped identify and quantify proteins and their modifications, offering insights into disease mechanisms. In an attempt to identify key players in breast cancer-induced muscle weakness, emerging studies have performed RNA sequencing and multiple omics analysis on skeletal muscle in mice ^32,33^ and cell culture^34^. We have recently reported that experimental lung cancer can cause insulin resistance in skeletal muscle and adipose tissue in mice^35–37^, suggesting that tumours can secrete factors that induce insulin resistance in the skeletal muscle. Concomitantly, our recent meta-analysis showed that patients with various cancers are markedly insulin-resistant^38^, illustrating an intriguing endocrine cross-talk between cancer and muscle cells. Yet, whether breast cancer induces alterations in the proteomics of healthy skeletal muscle and alters insulin action is unknown.

To address that knowledge gap, we incubated muscle cells with media conditioned by breast cancer cells to explore tumour-muscle crosstalk. We found that incubation with media from two different breast cancer cell lines induced proteomic changes in pathways related to sarcomere organisation, actin filament binding, and vesicle trafficking, disrupted the myogenic differentiation and activated an inflammatory response, together with impaired muscle GLUT4 translocation and glucose uptake. These changes were associated with reduced insulin-mediated Rac1, but not Akt-TBC1D4, signalling. Thus, we obtain evidence for the direct effects of breast cancer on skeletal muscle proteome and glucose metabolism regulation.

## Methods

GLUT4myc-L6 skeletal muscle cells. Rat L6 skeletal myoblasts stably over-expressing c-myc epitope-tagged GLUT4 (GLUT4myc-L6), a kind gift from Amira Klip^39^, were grown in α-MEM media (GIBCO #22571) with 10% fetal bovine serum (FBS) (GIBCO #23140), 1% penicillin (10,000 U/ml)/streptomycin (10 mg/ml)/Amphotericin B (25μg/ml) (P/S/A) (GIBCO #A5955). L6 myoblasts were differentiated into myotubes for 7 days in α-MEM medium with 2% FBS, and 1% P/S/A. Cells were cultured in a humidified incubator at 5% CO2 and 37 °C for all cell experiments.

### Breast cancer cell lines

The human female breast cancer cell lines, MCF7 (ER+, PR+, HER2-) and BT474 (ER+, PR+, HER2+), were purchased from the American Type Culture Collection (ATCC), Manassas, VA. MCF7 was cultured in Roswell Park Memorial Institute 1640 medium (RPMI, GIBCO #61870-010) supplemented with 6% FBS (5% CO_2_, 37°C). BT474 was cultured in Dulbecco’s modified Eagle’s medium and F12 medium (DMEM/F12, GIBCO #31330-038) supplemented with 10% FBS (5% CO_2_, 37°C). 1% P/S/A was added to all culture media. Cells were used for experiments within the same 10-15 passages, and they were authenticated by routine visual inspection of the morphologic phenotype.

### MCF10A breast epithelial cell line

Control non-tumorigenic semi-normal breast epithelial cell line, MCF10A^40^, was purchased from ATCC. MCF10A was cultured in DMEM/F12 supplemented with 5% Horse Serum (Gibco #26050070), 20ng/ml Epidermal Growth Factor (EGF, Sigma #E-4127), 0.5μg/ml Hydrocortisone (VWR # CALB3867), 100ng/ml Choleratoxin (Sigma #C-8052), 10μg/ml Insulin (Sigma #I9278) and 1% P/S/A (5% CO_2_, 37°C).

### Breast cancer-conditioned media preparation and treatment

Tumorigenic (MCF7 and BT474) breast cells were cultured in their respective growth media in a 100 mm dish (8*10^5^-10^6^ cells/dish in 10 ml growth media) reaching 80-90 % confluence after 48 hr. The growth media of the control non-tumorigenic (MCF10A) breast cells contains supplements that disturb the differentiation of GLUT4myc-L6 myotubes. Thus, the MCF10A conditioned media was prepared using Dulbecco’s Modified Eagle Medium (DMEM, Gibco # 11995073) with 10% FBS and 1% P/S/A. Here, the cells were seeded (10^6^ cells/dish) four days before use and allowed to acclimate for two days. After two days, the DMEM media was replenished (10%FBS, 1%P/S/A). After 48 hr, the media was removed from all breast cell lines, and centrifuged (1000 g, 5 min, room temperature (RT)). The supernatant, which is referred to as supernatants were diluted in fresh GLUT4myc-L6 differentiation media (40% conditioned media: 60% differentiation media), and used within 30 min to avoid protein denaturing and loss of activity of the tumour-secreted factors.

The 40% conditioned media was added on GLUT4myc-L6 myotubes that were already differentiated for 4 days in normal differentiation media. Myotubes were replenished with freshly prepared conditioned media every 24 hr over three days, resulting in 72 hours breast cancer-conditioned media treatment. Experiments were performed on day 7 on differentiated GLUT4myc-L6 myotubes.

### GLUT4 translocation

On the experimental day, GLUT4myc-L6 myotubes (day 7) were deprived of serum 3 hours before insulin stimulation (15 min stimulation, 10 nM insulin (Actrapid; Novo Nordisk, DK)), cells were stimulated for. At the end of the stimulation period, cells were chilled on ice, washed 3 times with ice-cold PBS and fixed in 3% paraformaldehyde (PFA) (10 min on ice, and 10 min in RT), blocked in 5% goat serum (GS), and incubated with a primary anti-c-Myc antibody (Sigma #C3956). The signal was detected with an HRP-conjugated anti-rabbit secondary antibody, and o-Phenylenediamine (OPD) reagent (Sigma #P5412) was added to each well to initiate a colour reaction with the secondary antibody. The reaction was terminated with 5N HCl and absorbance was measured at 492 nm. Background absorbance (no primary antibody) was subtracted from the measured values of absorbance. The experiment was assayed in n = 4-8 for each group and repeated 6 times.

### ^3^H-labeled 2-Deoxy-Glucose uptake in GLUT4myc-L6 cells

Glucose uptake in GLUT4myc-L6 myotubes was measured using the radio-labeled D-2-deoxy-D-[2,6-^3^H]glucose ([^3^H]-2DG) as described previously [27]. Breast cancer-conditioned media treated GLUT4myc-L6 myotubes (day 7) were incubated in serum-free medium for 3 hrs, and stimulated with human insulin (10 nM, Actrapid; Novo Nordisk, DK) for 20 min before the end of the starvation period, followed by 3X wash with HEPES buffered solution (140 mM NaCl, 20 mM HEPES, 5 mM KCl, 2.5 mM MgSO_4_, 1.0 mM CaCl_2_, pH 7.4 at 37°C) and further 5 min incubation with transport solution (10 µM 2DG and 10 µM ^3^H-2DG (1.0 µCi/well)) with or without insulin (10 nM). The 2DG transport was terminated by 3X wash with Stop solution (0.9% NaCl, 25 mM glucose). To assess non-specific glucose uptake, cells were treated with cytochalasin B (20 µM, Sigma #C6762). Cells were lyzed in 70 μl lysis buffer (10% Glycerol, 20 mM Na-pyrophosphate, 150 mM NaCl, 50 mM HEPES, 1% NP-40, 20 mM β-glycerophosphate, 1 M NaF, 2 mM PMSF, 1 mM EDTA, 1 mM EGTA, 10 µg/ml Aprotonin, 10 µg/ml Leupeptin, 2 mM Na_3_VO_4_, 3 mM Benzamidine). Lysates were clarified by centrifugation (10 min, 4 °C, 9000 *g*), and the supernatant was collected and stored at −80 °C. To measure specific glucose uptake, lysates were assessed for [^3^H]-2DG tracer counts by scintillation counting using a β-scintillation counter. All tracer counts were normalized to total protein content. The experiment was repeated 5 times with n = 3-5.

### Cell lysate

Cell lysates were prepared from GLUT4myc-L6 myotubes (day 7) incubated with conditioned media from MCF7, BT474, and MCF10A breast cells as described above (see *^3^H-labeled 2-Deoxy-Glucose uptake in GLUT4myc-L6 cells*). Protein concentrations were determined by the bicinchoninic acid (BCA) method using BSA standards and BCA assay reagents (Pierce).

### Immunoblotting

Total protein and phosphorylation levels of relevant proteins were determined by standard immunoblotting technique loading 4-6 µg of protein to investigate intracellular signalling in response to insulin. The polyvinylidene difluoride membrane (Immobilon Transfer Membrane; Millipore) was blocked in TBS-Tween 20 containing 2% skim milk for 15 minutes at room temperature, followed by incubation overnight at 4°C with the primary antibody. The primary antibodies used were Akt2 (#3063, Cell Signalling Technology (CST)), p-Akt S473 (#4058, CST), p-Akt T308 (#4058, CST), TBC1D4 (ab189890, Abcam), p-TBC1D4 T642 (#D27E6, CST), RhoGDIα (#2564, CST), Rac1 (#610651, BD bioscience), Cofilin-2 (#sc-166958, Santa Cruz), p-Cofilin S3 (#3313, CST), and GAPDH (#2118, CST). Next, the membrane was incubated with HRP-conjugated secondary antibody (Jackson Immuno Research) for 45-60 min at room temperature. Bands were visualized using Bio-Rad ChemiDocTM MP Imaging System and enhanced chemiluminescence (ECL+; Amersham Biosciences). Reversible Protein Stain Kit (Pierce #24585) was used as a loading control.

### Rac1 activity assay

Rac1 activity (GTP bound Rac1) was measured using a commercially available colorimetric Rac1 G-LISA Activation Assay kit (Cytoskeleton Inc. # BK126) following the manufacturer’s instructions. Briefly, after conditioned media treatment, cells underwent 3 hr serum starvation and 10 min of insulin stimulation (10 nM). After the treatment, the cells were quickly lysed in ice-cold lysis buffer containing a protease inhibitor cocktail, and the samples were clarified by centrifugation (8792 *g*, 1 min, 4 °C). The supernatant was collected and snap-frozen in liquid nitrogen for storage at −80 °C. Using the BCA assay described earlier, the protein concentration was measured and 30 µg protein was loaded into a 96-well plate coated with Rac1 binding domains. The plate was incubated on a shaker (30 min, 4 °C) and the active GTP-bound Rac1 was determined colourimetrically by specific Rac1 antibodies, and absorbance was measured at 492 nm.

### Immunofluorescence

GLUT4myc-L6 cells were plated in µ-slide 8-well polymer coverslip bottom (#808806, Ibidi) plates. Myotubes were fixed in 4% PFA, permeabilised in PBS containing 0.2% Triton X for 10 min and blocked in 5% GS for 45 min. Cells were then incubated with primary antibody against Myosin Heavy Chain (MHC) (1:50, #MF20 IgG2b, Developmental Studies Hybridoma Bank, USA) and α-Tubulin (1:500, #IgG1, T9026, Sigma) at 4°C overnight in 5% GS. The fluorescence was detected using the secondary antibody red-fluorescent Alexa-flour 568 (1:1000, IgG2b, #A-21144, Invitrogen, 1 hr RT) and magenta-fluorescent Alexa-flour 647 goat-anti-mouse (1:500, #A-21235, Invitrogen, 1 hr, RT). To stain against actin filaments, cells were incubated in green-fluorescent Alexa-flour 488 Phalloidin (1:400, #8878, Cell signalling, 1hr RT). Cells were then washed 3×3 min, with one wash including DAPI (1:50000, #D3571, Invitrogen) to stain for nuclei, and cells were mounted in Prolong Gold and stored at 4°C. For negative controls, 1-2 wells were not added primary antibody to ensure specificity.

### Ruffling index quantification

All fluorescence microscopy was performed with Zeiss Cell Observer spinning disc confocal microscopy using a 63x, 1.4 numerical aperture oil objective. Images were acquired in 0.45 µm *z*-increments. This high-resolution objective was chosen to enable clear delineation between cell borders and to distinguish between myotubes and myoblasts within each *z*-plane. Cells were analysed in 3D using FIJI (insert ref) to determine ruffles based on cell morphology and actin intensity. The ruffling index was quantified by identifying F-actin-rich regions extending to the dorsal plane^41^, and α-tubulin staining was used to confirm/verify membrane ruffling. A total of 80-200 myotubes and 800-900 myoblasts were analysed from three separate experiments with n=3 per condition. Ruffling index quantifications are presented as the average number of ruffles per cell.

### Myotube width measurements

To quantify myotube width, Tubulin, MHC and phalloidin stains were used to determine myotube borders. The average of three measurements of each myotube was taken. 9-14 myotubes were analyzed in each condition.

### RNA extraction and real-time-qPCR

RNA extraction from GLUT4myc-L6 myotubes was conducted using RiboPure™ RNA Purification Kit (Invitrogen #AM1924) following the manufacturer’s instructions. RNA was isolated by using TRI Reagent and subsequently purified using Filter Cartridges. The isolated RNA was reverse transcripted using Q-Script cDNA SuperMix (QuantaBio), and PCR was performed by LightCycler LC480 Real-Time PCR System (Roche Applied Science) and the SYBR Green PCR MasterMix (Applied Biosystems). All measurements were related to *sdha*. All predesigned primers for rat species (KiCqStart® Primers, Sigma, DK) are listed in Table 1.

**Table 1.**
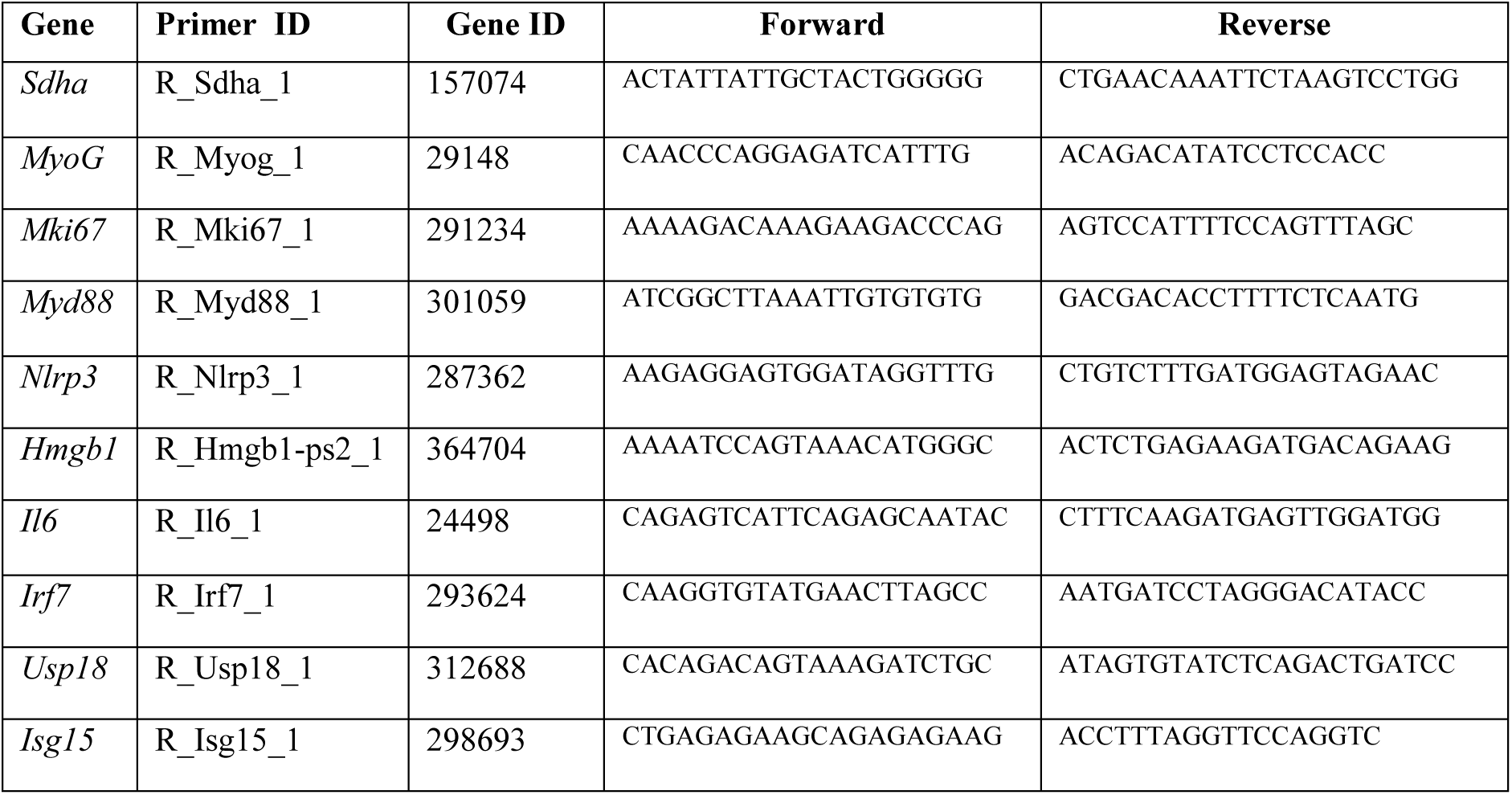
List of predesigned primers for rat species used in the current study (KiCqStart® Primers, Sigma, DK).

### Cell pellet preparation for Proteomics

GLUT4myc-L6 myotubes underwent a 3-hour fasting period in serum-free α-MEM medium. Cells were then washed twice with warm PBS and treated with trypsin (Gibco, #25200056) for 2 minutes at 37°C. The detached cells were collected by adding growth medium to the cell culture plates and resuspending the cells. Next, the cell suspension was centrifuged (1500 rpm, 2 min, RT), and the supernatant was discarded. The cell pellet was resuspended in ice-cold PBS, and centrifuged (2000 rpm, 2 min, 4°C), followed by an additional wash in ice-cold PBS and centrifugation. Finally, all PBS was removed from the pellet, and the samples were snap-frozen in liquid nitrogen and stored at −80°C for later analysis.

### Protein digestion and Tandem Mass Tags (TMT) labelling

Cell pellets were prepared on the AccelerOme robot (Thermo Fisher Scientific) using the AcclerOme TMTpro 16plex MS Sample Preparation and Lanelign Kit (A50950). Briefly, cells were lysed in lysis buffer in the presence of nuclease by pipetting and incubated for 10 min at room temperature. Lysates were cleared by centrifugation (16,000g for 5 min) and supernatants were sonicated in a Bioruptor Pico for 5 min. Samples were incubated at 95 °C for 10 min, and proteins measured and adjusted in lysis buffer, and digested into peptides before processing with the AccelerOme for TMT labelling.

### High-pH fractionation

Labelled desalted peptides were resuspended in buffer A* (5% acetonitrile, 1% TFA), and fractionated into 16 fractions by high-pH fractionation. For this, 20 ug peptides were loaded onto a Kinetex 2.6u EVO C18 100 Å 150 x 0.3 mm column via an EASY-nLC 1200 HPLC (Thermo Fisher Scientific) in buffer AF (10 mM TEAB), and separated with a non-linear gradient of 5 – 44 % buffer BF (10mM TEAB, 80 % acetonitrile) at a flow rate of 1.5 µL / min over 62 min. Fractions were collected every 60 s with a concatenation scheme to reach 16 final fractions (e.g. fraction 17 was collected together with fraction 1, fraction 18 together with fraction 2, and so on).

### Liquid Chromatography-Mass Spectrometry (LC-MS/MS)

The fractions were evaporated, resuspended in buffer A*, and measured on a Vanquish Neo HPLC system (Thermo Fisher Scientific) coupled through a nano-electrospray source to a Tribrid Ascend mass spectrometer (Thermo Fisher Scientific). Peptides were loaded in buffer A (0.1 % formic acid) onto a 110 cm μPAC HPLC column (Thermo Fisher Scientific) and separated with a non-linear gradient of 1 – 50 % buffer B (0.1 % formic acid, 80 % acetonitrile) at a flow rate of 300 nl/min over 100 min. The column temperature was kept at 50° C. Samples were acquired using a Real Time Search (RTS) MS3 data acquisition where the Tribrid mass spectrometer was switching between a full scan (120 K resolution, 50 ms max. injection time, AGC target 100%) in the Orbitrap analyzer, to a data-dependent MS/MS scans in the Ion Trap analyzer (Turbo scan rate, 23 ms max. injection time, AGC target 100% and HCD activation type). The isolation window was set to 0.5 (m/z), and normalized collision energy to 32. Precursors were filtered by a charge state of 2-5 and multiple sequencing of peptides was minimized by excluding the selected peptide candidates for 60 s.

MS/MS spectra were searched in real time on the instrument control computer using the Comet search engine (UniProtKB #UP000002494 Rattus Norvegicus FASTA file, 0 max miss cleavage, 1 max oxidation on methionine as variable mod. and 35ms max search time with an Xcorr scoring threshold of 1.4 and 20 precursor ppm error). MS/MS spectra resulting in a positive RTS identification were further analyzed in MS3 mode using the Orbitrap analyzer (45k resolution, 105 ms max. injection time, AGC target 500%, HCD collision energy 55 and SPS = 10). The total fixed cycle time, switching between all 3 MS scan types, was set to 3 s.

### Data analysis

Raw mass spectrometry data were analysed with Proteome Discoverer (v3.1). “PWF_Tribrid_TMTpro_SPS_MS3_SequestHT_INFERYS_Rescoring_Percolator” was used as default: processing workflow. Briefly, peak lists were searched against the UniProtKB #UP000002494 Rattus Norvegicus FASTA database by the integrated SequestHT search engine, setting Carbamidomethyl (C) and TMTpro (K, N-Term) as static modifications, Oxidation (M) as variable modification, max missed cleavage as 2 and minimum peptide amino acid length as 7. The false discovery rate was set to 0.01 (strict) and 0.05 (relaxed).

### Data availability

The mass spectrometry proteomics data have been deposited to the ProteomeXchange Consortium (http://proteomecentral.proteomexchange.org) via the PRIDE partner repository with the data set identifier PXDxxxxxx (https://www.ebi.ac.uk/pride/archive/projects/PXDxxxxxx).

### Bioinformatics

All statistical analysis of protein expression intensity data was conducted using custom Python 3.9.0 code (Proteomelit, version 1.0.0), and packages pandas (v2.0.1), scipy (v1.11.2), scikit-learn (v1.2.2), statsmodels (v0.14.0) and plotly (5.14.1). Data was searched against the rat (Rattus norvegicus) UniProt database (July 2024). Protein intensities were log2-transformed. Proteins with fewer than two valid values in at least one experimental group were excluded from the analysis. Missing values were estimated using the MinProb approach (width = 0.3, shift = 1.8) ^42^. MinProb method was used for values missing not at random (MNAR), defined when at least 60% of the samples within a given experimental group have a valid value. Differentially regulated proteins were identified using unpaired t-tests, and the *p-values* were adjusted for multiple hypothesis testing using two distinct methods: Benjamini-Hochberg (BH) correction or permutation-based false discovery rate (FDR) correction ^43^. Both tests included a false discovery rate of 5%. Benjamini-Hochberg method was applied with an FDR of 5%, and a fold change (FC) cut-off of 2. Permutation-based FDR correction was applied with an FDR of 5%, an FC cut-off of 1, s0 of 1, and 250 permutations. Permutation-based corrected *p-values* < 0.05 were considered significant and used in the enrichment analysis. Comparative analyses were conducted between MCF7 or BT474 conditioned media-treated myotubes and MCF10A conditioned media-treated myotubes. Differentially regulated proteins are visualized in upset plots and highlighted in volcano plots.

### Gene ontology (GO) enrichment analysis

Functional enrichment analysis was carried out using Fisher’s exact test, followed by Benjamini-Hochberg correction. Enrichment analysis included GO Biological Process, GO Cellular Component, and GO Molecular Function annotations. All proteins quantified in the study served as the background, while only statistically significant proteins were considered as the foreground. Enrichment analyses were also conducted on all overlapping regulated proteins between MCF7 and BT474 conditioned media compared to MCF10A conditioned media. Enrichment plots display only significant terms (adjusted p-value < 0.05) that were associated with at least two proteins from the foreground.

### Statistical analyses

Data are shown in superplots^44^ with mean ± SEM including individual variables that are shape- and color-coded according to their replicates. Statistical significance was tested using one-way ANOVA, or two-way ANOVA followed by Sidak’s post hoc test to evaluate the main effects and significant interactions. Separate two-way ANOVAs were performed for each breast cancer conditioned media compared to control to better identify the interaction specific to each breast cancer cell line. Individual values were used for statistical analyses. If a significant difference between experimental replicas was detected, the analysis was performed on the mean of each experimental round. The significance level was set at *p < 0.05*, and *p<0.1* was considered a trend. Statistical analyses were performed using Graph Pad PRISM v.9 software (GraphPad Software, Inc., San Diego, CA).

## Results

### Breast cancer-secreted factors induce proteomic changes related to sarcomere assembly, actin cytoskeleton remodelling and vesicle trafficking

To first establish whether breast cancer-secreted factors could directly alter the global skeletal muscle proteome, we performed mass spectrometry on healthy, insulin-sensitive GLUT4myc-L6 myotubes treated with conditioned media from MCF7 and BT474 breast cancer cells and non-tumourigenic MCF10A breast cells (Fig. 1A). To our knowledge, the direct effect of luminal A and B breast cancer-secreted factors on the global proteome of healthy myotubes has not been previously investigated. The conditioned media contains a large accumulation of secreted signalling molecules that are unique to each cell line ^45,46^. Thus, the secreted factors from the tumorigenic cells mimic the tumour microenvironment, which would not be present in the non-tumourigenic MCF10A breast cell-conditioned media. We hypothesized that the breast cancer-secreted factors in the conditioned media would directly influence the skeletal muscle. Mass spectrometry-based proteomic analysis revealed 259 and 122 regulated proteins in myotubes treated with MCF7 and BT474 conditioned media, respectively, compared to control MCF10A conditioned media (Fig. 1B). Of these, 63 regulated proteins were shared between myotubes exposed to MCF7 and BT474 conditioned media, of which 26 were upregulated, and 37 were downregulated (Fig. 1B). Thus, breast cancer-secreted factors induced alterations in the myotubes proteome.

**Figure 1.**
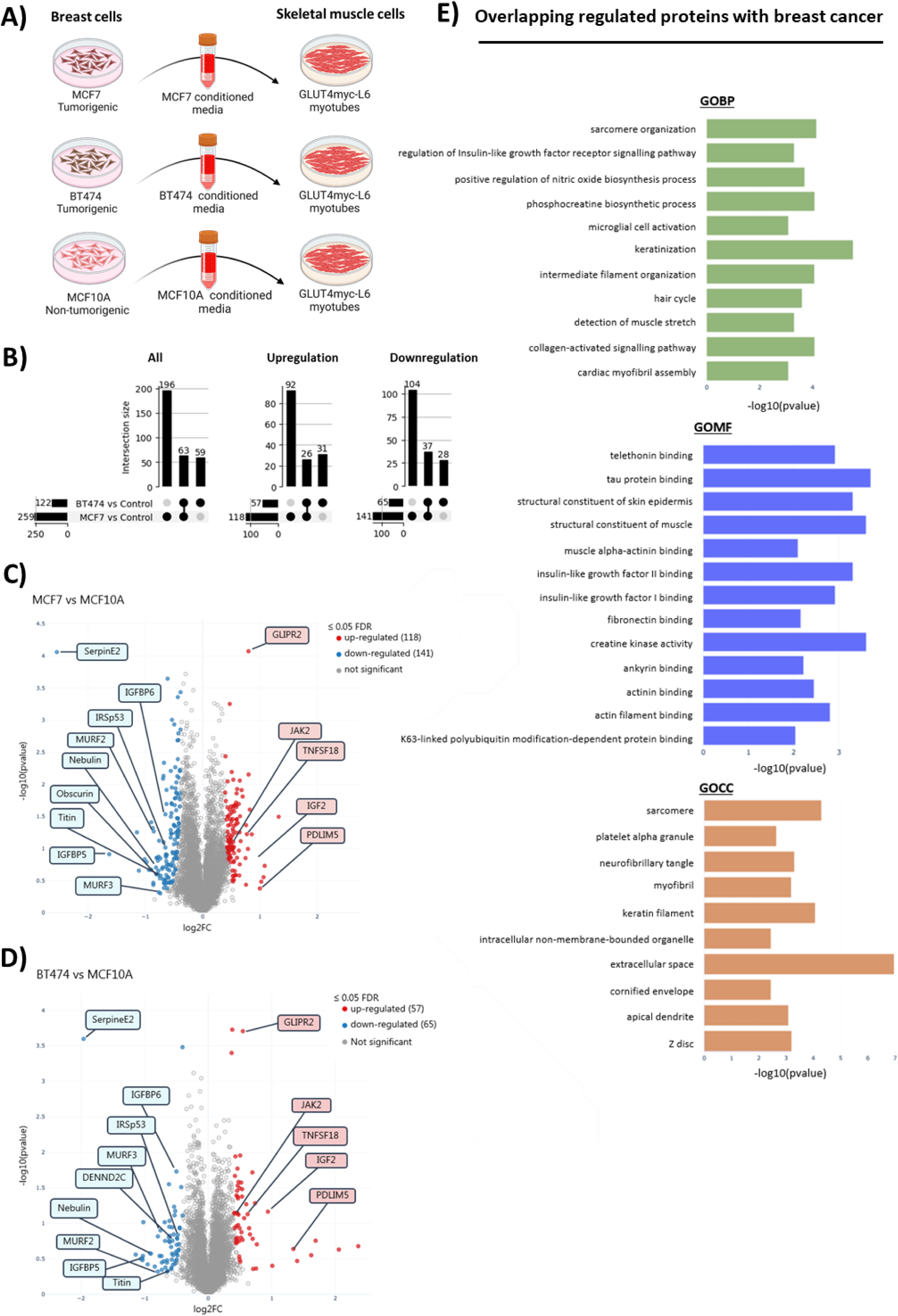
Breast cancer-secreted factors induce proteomic changes related to sarcomere assembly, actin cytoskeleton remodelling and vesicle trafficking. Mass spectrometry-based proteomic analysis of GLUT4myc-L6 myotubes treated with MCF7 and BT474 breast cancer-conditioned media treatment compared to control MCF10A media. N=3. **A)** Schematic illustration of the experimental workflow. GLUT4myc-L6 myoblasts were differentiated into myotubes for 4 days in regular L6 differentiation media. When GLUT4myc-L6 myotubes are differentiated for 2 days, the breast cells were seeded accordingly to reach 80-90% confluency after 48hr. After 48 hours, the conditioned media (CM) was removed from the breast cells, centrifuged (800-1000 *g*, 5 min, RT) to remove dead cells and debris, and the supernatants were collected. A 40% dilution of the conditioned media in L6 differentiation media was immediately added to the GLUT4myc-L6 myotubes (day 4). The conditioned media on GLUT4myc-L6 myotubes was replenished every 24 hr for three days, resulting in 72 hours breast cancer-conditioned media treatment. Created with biorender.com. **B)** Up-set plot of differently regulated proteins (Significance score ≤0.05; permutation-based FDR correction with an FDR of 5%). Volcano plot comparing protein abundance induced by **C)** MCF7 and **D)** BT474 conditioned media treatment compared to control MCF10A media. Differently regulated proteins are marked in blue (upregulation) and red (downregulation). **E)** Fischer’s exact test for enrichment analysis of the 63 overlapping regulated proteins (Significance score ≤0.05; Benjamini-Hochberg correction with an FDR of 5%). Gene ontology molecular function (GOMF), cellular component (GOCC) and biological processes (GOBP). Data was searched against the rat (Rattus norvegicus) UniProt database.

Roughly 50% of the overlapping regulated proteins were implicated in myogenic differentiation and sarcomere organization. The sarcomere is a key indicator of myotube maturation, which provides structural stability and alignment into the muscle cell. Figure 1C and 1D show volcano plots with selected proteins highlighted, which are all involved in pathways related to sarcomere, actin cytoskeleton binding, and myogenic differentiation (Fig. 1C-E). We observed downregulation in several sarcomeric component proteins including Titin, Obscurin, and Nebulin as well as Trim55/54 (known as MURF2/3) ^47^. Interestingly, breast cancer-conditioned media upregulated several LIM domain proteins, including CSRP3, PDLIM5, and GLIPR2^48^, which are implicated in muscle mass regulation and myofiber organization via autophagy^49^. Overexpression of PDLIM5 is shown to increase proliferation and inhibit cell cycle exit in skeletal muscle cells^50^. Notably, breast cancer cell-conditioned media downregulated the protease inhibitor, SERPINE2, known as Protease Nexin-1 (PN1), which promotes myoblast-fusion into multinucleated myotubes^51^. These findings suggest that breast cancer-secreted factors reduce the ability of myoblasts to exit the cell cycle and fuse into multinucleated myotubes, hence, reducing myogenic differentiation.

Additionally, breast cancer cell-conditioned media deregulated several proteins involved in the actin cytoskeletal remodelling (Fig. 1D). Importantly, we observed downregulation in proteins involved in Rho GTPase signalling including BAIAP2 (also known as IRSp53)^52^ and ABRA (Actin binding Rho activating protein, also known as STARS)^53^. IRSp53 is especially of interest to this study. Although not confirmed in muscle, IRSp53 has been suggested to serve as a Rac1 downstream effector protein^52^, inducing actin cytoskeletal remodelling, which is important for insulin-stimulated glucose uptake and vesicle trafficking^23–27,54,55^. Moreover, breast cancer cell-conditioned media led to downregulation in DENND2C, a known activator of Rab GTPases^56^, which drives vesicle exocytosis, a crucial step for glucose uptake in response to insulin^57–59^. These results suggest that breast cancer-secreted factors deregulate important proteins involved in insulin-stimulated glucose uptake.

Furthermore, breast cancer-conditioned media upregulated several proteins involved in pro-inflammatory pathways related to muscle atrophy^60^. Among these, TNFSF18 (TNF superfamily 18)^61^, ASAH1 (Acid ceramidase), PTGS2 (known as COX2), and JAK2 (Janus Kinase 2) were upregulated. JAK2 is involved in the JAK/STAT pathway, activated by IL6 through STAT3^62^, promoting myogenic differentiation via IGF2^63–65^. Interestingly, we observed an increased level of IGF2 protein with breast cancer-conditioned media, which could increase IL6 expression by activating NF-ĸB pathway as shown in immune ^66^ and muscle cells^67^. On the other hand, the IGF binding protein-5 and -6 (IGFBP-5/6) were downregulated. The absence of the muscle-specific IGFBP-5 has been shown to reduce the expression of the myotube-marker myosin heavy chain (MHC)^68^. Together, these results suggest that breast cancer-secreted factors directly alter the proteome of myotubes in pathways related to sarcomere organization, myogenic programming, actin cytoskeleton remodelling, vesicle trafficking, and inflammation.

### Breast cancer-conditioned media decreases differentiation and induces inflammation of GLUT4myc-L6 myotubes

Previous studies have documented increased inflammation and muscle dysfunction in association with breast cancer both *in vitro*^34,69^ and *in vivo*^29,31–33^, aligning with our observations from proteomics analyses. Therefore, we investigated whether breast cancer-secreted factors from MCF7 and BT474 cells directly affect the myogenic differentiation in GLUT4myc-L6 myotubes. During myogenic differentiation, myoblasts exit the cell cycle and fuse into elongated and multinucleated myotubes. Several myogenic regulatory factors are involved in each step of this process including myogenin (*MyoG*)^70,71^. We found no effect of breast cancer treatment on *MyoG* mRNA expression (Fig. 2A). On the other hand, the proliferation marker *Mki67* (encoding Ki67 protein) was elevated by 50% (*p=0.009*) with breast cancer-conditioned media compared to control MCF10A conditioned media (Fig. 2A). These data suggest that breast cancer affects skeletal muscle myogenesis by preventing myoblasts from exiting the cell cycle, which is supported by our proteomics data. To confirm whether breast cancer cell-conditioned media influences the differentiation of GLUT4myc-L6 myoblasts, we measured MHC expression by immunofluorescence staining. Treatment with conditioned media from both MCF7 and BT474 breast cancer cells markedly reduced MHC expression in GLUT4myc-L6 cells compared to MCF10A conditioned media (Fig. 2B).

**Figure 2.**
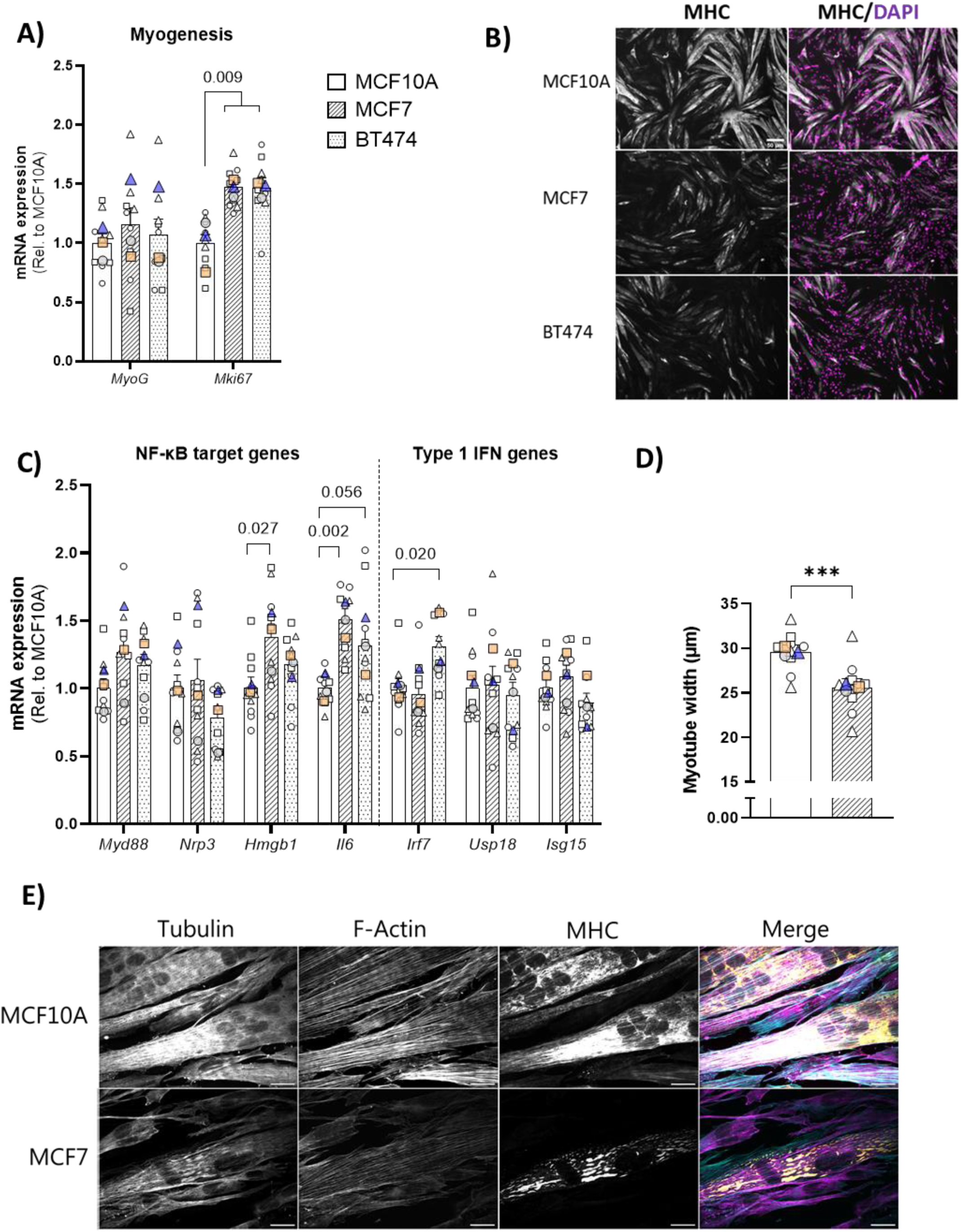
Breast cancer-conditioned media decreases differentiation and induces inflammation of GLUT4myc-L6 myotubes. **A)** Myogenesis target genes in GLUT4myc-L6 myotubes treated with 40% MCF7 (striped), BT474 (dotted) and MCF10A (plain) conditioned media for 72 hr. N=3 repeated three times. **B)** Representative immunofluorescence images of GLUT4myc-L6 myotubes incubated with 40% conditioned media from MCF7, BT474, and MCF10A during basal state using co-staining for MHC (Gray), and DAPI (Magneta; nuclear stain). Scalebar = 50 µm at 4X magnification. N=2 repeated twice. **C)** NF-κB target and type I IFN response gene expression in GLUT4myc-L6 myotubes treated with 40% MCF7 (striped), BT474 (dotted) and MCF10A (plain) conditioned media for 72 hr. N=3 repeated three times. **D)** Myotube width in (µm) measured on myotubes treated with 40% MCF10A and MCF7 conditioned media. N=3 repeated three times. **E)** Representative immunofluorescence images of GLUT4myc-L6 myotubes incubated with 40% conditioned media from MCF7 and MCF10A at baseline using co-staining for α-Tubule (Gray-scale/Magenta), F-Actin (Gray-scale/turquoise), and myosin heavy chain (Gray-scale/Yellow). Scalebar = 200 µm at 63X magnification. N=3 repeated thrice. Data in **A)** and **C)** are presented in super plots as mean ±SEM incl. individual values that are symbol-coded (representing wells) matching each experimental round (large colour-/and shape-coded symbols representing the average of each experimental round)^44^. All data in **A)** and **C)** were evaluated with one-way ANOVA using pool of individual values from all three round, except for gene *Mki67* and *Irf7* where mean values were used. Significant multiple comparisons in the ANOVAs were evaluated by Sidak’s post hoc tests: The exact p-values of breast cancer-conditioned media effect are stated on the graph when above p>0.001.

Besides myogenic regulatory factors, other signalling pathways also regulate skeletal muscle differentiation, including the nuclear factor-κB (NF-κB) and Type 1 interferon (IFN) inflammatory pathways^72–74^. MCF7 conditioned media treatment led to an upregulation of the NF-κB target genes *Hmgb1*^75^ and *Il6*. BT474 conditioned media increased NF-κB-related *Il6* gene expression, and type 1 IFN-related target gene *Irf7*^76^ compared to control MCF10A media (Fig. 2C). These results suggest that breast cancer-secreted factors trigger an intracellular pro-inflammatory response through NF-κB and type 1 IFN pathway, disrupting skeletal muscle myogenic programming.

Besides affecting myogenic differentiation, activation of the type 1 IFN and NF-κB pathways is also implicated in muscle atrophy^77,78^. Given the more apparent activation of the NF-κB pathway with MCF7 conditioned media, we chose to measure myotube width in response to MCF7 conditioned media treatment. Expectedly, myotube width was decreased by 15% with MCF7 conditioned media (Fig. 2D). These results suggest that breast cancer-secreted factors disrupt the myogenic differentiation by inducing muscle atrophy through activation of the pro-inflammatory NF-κB pathway. Furthermore, we observed concomitant MHC fragmentation, a potential marker for muscle atrophy ^79^, in myotubes treated with MCF7 conditioned media (Fig. 2E). This observation is consistent with the disrupted sarcomere organization in the proteomic analysis. Altogether, these findings suggest that breast cancer-secreted factors induce muscle atrophy likely via activation of the pro-inflammatory NF-κB.

### Breast cancer-derived factors inhibit skeletal muscle basal and insulin-stimulated GLUT4 translocation and glucose uptake

Given that NF-κB overexpression reduces glucose uptake and GLUT4 translocation in skeletal muscle cells^77^, we hypothesized whether the conditioned media from breast cancer cells would disrupt insulin sensitivity in myotubes. This is corroborated by our results from the proteomics analysis showing deregulated proteins involved in actin cytoskeleton remodelling and vesicle trafficking (Figure 1C,D), which are two key processes in regulating insulin-stimulated glucose uptake. A key step in skeletal muscle glucose uptake is the ability of glucose transporter, GLUT4, to translocate to the plasma membrane^80^ (Fig. 3A). Accordingly, lower GLUT4 translocation and impaired insulin-stimulated glucose uptake have been detected in multiple insulin-resistant conditions such as type 2 diabetes ^20,81^, obesity ^82,83^, and cancer ^35–38^. We observed reduced basal GLUT4 translocation in GLUT4myc-L6 myotubes incubated with conditioned media from MCF7 (−35%, *p < 0.001)* and BT474 (−20%, *p < 0.001*) compared to the control MCF10A conditioned media-treated muscle cells (Fig. 3B). The insulin-stimulated GLUT4 translocation was reduced by conditioned media from MCF7 (−20%, *p < 0.001)*, and BT474 conditioned media (−10%, *p < 0.001*) compared to control MCF10A conditioned media (Fig. 3B). These findings indicate that exposure to breast cancer-derived factors inhibit skeletal muscle basal and insulin-stimulated GLUT4 translocation.

**Figure 3.**
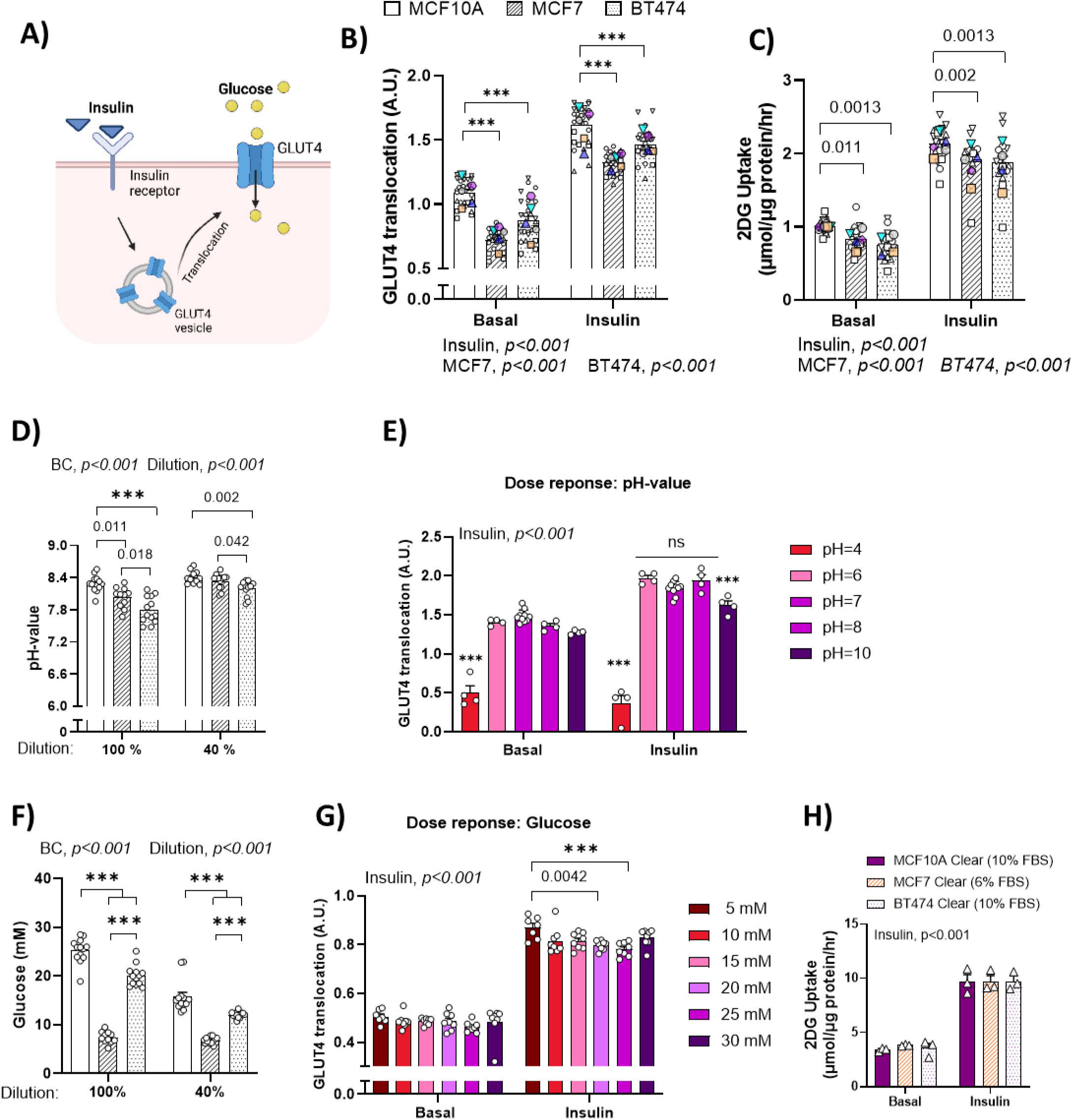
Breast cancer-derived factors inhibit skeletal muscle basal and insulin-stimulated GLUT4 translocation and glucose uptake. **A)** Schematic illustration of insulin-stimulated GLUT4 translocation and glucose uptake. Created with biorender.com. **B)** Insulin-stimulated (10 nM, 15 minutes) GLUT4 translocation in GLUT4myc-L6 myotubes incubated with 40 % conditioned media from tumorigenic MCF7 (striped bars) or BT474 (dotted bars) cells for 72 hrs compared to control non-tumorigenic MCF10A (plain bars) cells. N = 4-8 repeated six times. **C)** Basal and insulin-stimulated (10 nM, 25 minutes) 2-deoxyglucose (2DG) uptake in GLUT4myc-L6 myotubes incubated with 40 % conditioned media from MCF7 (striped bars) or BT474 (dotted bars) cells for 72 hrs compared to control MCF10A (plain bars) cells. N = 3-5 repeated five times. **D)** pH of the conditioned media from MCF7, BT474 and MCF10A cell lines, before (100%) and after 40% dilution in the differentiation medium. N=11-13. Data shown in ±SEM with individual values. **E)** pH dose-response on basal and insulin(10 nM, 15 min)-stimulated GLUT4 translocation in GLUT4myc-L6 myotubes. N=4-10. Data shown in ±SEM with individual values. **F)** Glucose concentrations (mM) of the conditioned media from MCF7, BT474 and MCF10A cell lines, before (100%) and after 40% dilution in the differentiation medium. N=11-13. Data shown in ±SEM with individual values. **G)** Glucose (mM) dose-response on basal and insulin (10 nM, 15 min)-stimulated GLUT4 translocation. N=7-8. **H)** Basal and insulin-stimulated (10 nM, 25 minutes) 2DG uptake in GLUT4myc-L6 myotubes incubated with 40% non-conditioned growth media for each breast cell line MCF7=RPMI (6% FBS, striped bars), BT474=DMEM/F12 (10%FBS, dotted bars), MCF10A=DMEM (10% FBS, plain bars) for 72 hrs. N = 3 repeated once. Data shown in ±SEM with individual values. For (B,C) data are presented in super-plots as mean ±SEM incl. individual values that are symbol-coded (representing wells) according to each round (large colour-/and shape-coded symbols representing the average of each experimental round)^44^. Data were evaluated with (B-D,F,H) two-way ANOVA using pool of individual values, and (E,G) mixed-effect analysis. Main effects of insulin, MCF7 or BT474 conditioned media treatment are indicated in the panels. Significant main effect and interaction in two-way ANOVAs were evaluated by Sidak’s post hoc: The exact p-values of MCF7 or BT474 conditioned media effect are stated on the graph when p>0.001 or *** if below p<0.001. A.U., arbitrary units

Having observed a reduction in GLUT4 translocation, we next determined basal and insulin-stimulated glucose uptake. In line with the reduction in GLUT4 translocation, MCF7 and BT474 conditioned media lowered basal glucose uptake by −20% (*p=0.011*) and −30% (*p=0.0113*), respectively (Fig. 3C). Insulin-stimulated glucose uptake was also reduced by −10% with both MCF7 (*p=0.002*) and BT474 (*p=0.0013*) conditioned media-incubation compared to control MCF10A conditioned media (Fig. 3C). Overall, these results suggest that secreted factors from breast cancer cells lower basal and insulin-stimulated skeletal muscle glucose uptake by reducing GLUT4 translocation to the plasma membrane.

To control for potential confounding factors in the conditioned media, we tested the pH and glucose concentrations of the conditioned media from all three cell lines, before and after 40% dilution in the differentiation medium. All three conditioned media had a pH of 7.8 to 8.3 before dilution, which increased (by 0.1-0.4 pH units) after dilution (Fig. 3D). Both MCF7 and BT474 conditioned media had a slightly lower (−0.2 and −0.5 units) pH than the MCF10A control media. Although the changes we observed were minor, we tested if media pH could affect basal or insulin-stimulated GLUT4 translocation, by performing a pH-dose response experiment (Fig. 3E). Insulin-stimulated GLUT4 translocation was slightly decreased at pH of 10, while the pH of 4 completely abrogated basal and insulin-mediated GLUT4 translocation. No changes were detected within the pH range of 6 to 8, which was our experimental range. Therefore, it is unlikely that the effects of breast cancer-conditioned media were caused by differences in pH levels.

Cancer cells are known to exhibit high levels of glucose consumption^84^, which could in turn influence the glucose concentration levels in the conditioned media. In line with that, the glucose levels in the media from the tumourigenic MCF7 and BT474 were lower than in the conditioned media from the control MCF10A (Fig. 3F). To examine the effect of glucose concentration on GLUT4 translocation, we performed a glucose dose-response experiment on GLUT4myc-L6 myotubes. Neither basal nor insulin-stimulated GLUT4 translocation was affected by glucose concentrations between 10 mM and 30 mM (Fig. 3G).

Several studies have demonstrated that serum *per se* can induce glucose uptake in myotubes independent of insulin ^85–87^. In our experiments, breast cancer cells were cultured in growth media containing different concentrations of FBS (MCF10A and BT474= 10% FBS, MCF7= 6% FBS). To assess whether these differences in FBS concentrations influenced glucose uptake, we incubated GLUT4myc-L6 myotubes with non-conditioned media that did not contain breast cancer-secreted factors. The non-conditioned media was prepared in the same ratio as the 40% breast cancer cell-conditioned media, consisting of a 2:5 ratio of the respective breast cell growth media (40%) to GLUT4myc-L6 differentiation media (60%). We found no effect of the different 40% non-conditioned media on basal or insulin-stimulated glucose uptake in GLUT4myc-L6 myotubes (Fig. 3H). Thus, any potential differences in pH, glucose concentrations, or FBS between the conditioned media do not influence the observed effects of breast cancer cell-conditioned media on GLUT4 translocation and glucose uptake. Altogether, these results show that breast cancer-derived factors inhibit skeletal muscle basal and insulin-stimulated GLUT4 translocation and glucose uptake in myotubes.

### Breast cancer-secreted factors increase myotube Akt2 and TBC1D4 protein content without altering insulin-stimulated Akt-TBC1D4 phosphorylation

Having established that conditioned media from breast cancer cells lowered GLUT4 translocation and glucose uptake, we next investigated the potential molecular mechanisms modulating these changes by analysing the intramyocellular insulin signalling (Fig. 4A). We observed no effect of MCF7 or BT474 conditioned media on phosphorylated (p)-Akt T308 and p-Akt S473 in the myotubes (Fig. 4B, C). In agreement, we found no difference in p-TBC1D4 T642 (a downstream target of Akt) by breast cancer-conditioned media (Fig. 4D). Interestingly, total Akt2 protein content was elevated (MCF7, 30% *p=0.005* and BT474, 40%, *p=0.002*) in myotubes treated with breast cancer-conditioned media (Fig. 4E). Similarly, TBC1D4 protein content was increased by MCF7 (20%, *p=0.020*) and BT474 (40%, *p=0.003*) (Fig. 4F). Thus, when relating phosphorylated Akt2 and TBC1D4 to total protein, there was a slight reduction in p-Akt T308 and S473, but not p-TBC1D4 T642. These data suggest that insulin signalling via Akt-TBC1D4 is not responsible for the impairments in basal and insulin-stimulated GLUT4 translocation and glucose uptake with breast cancer-conditioned media treatment.

**Figure 4.**
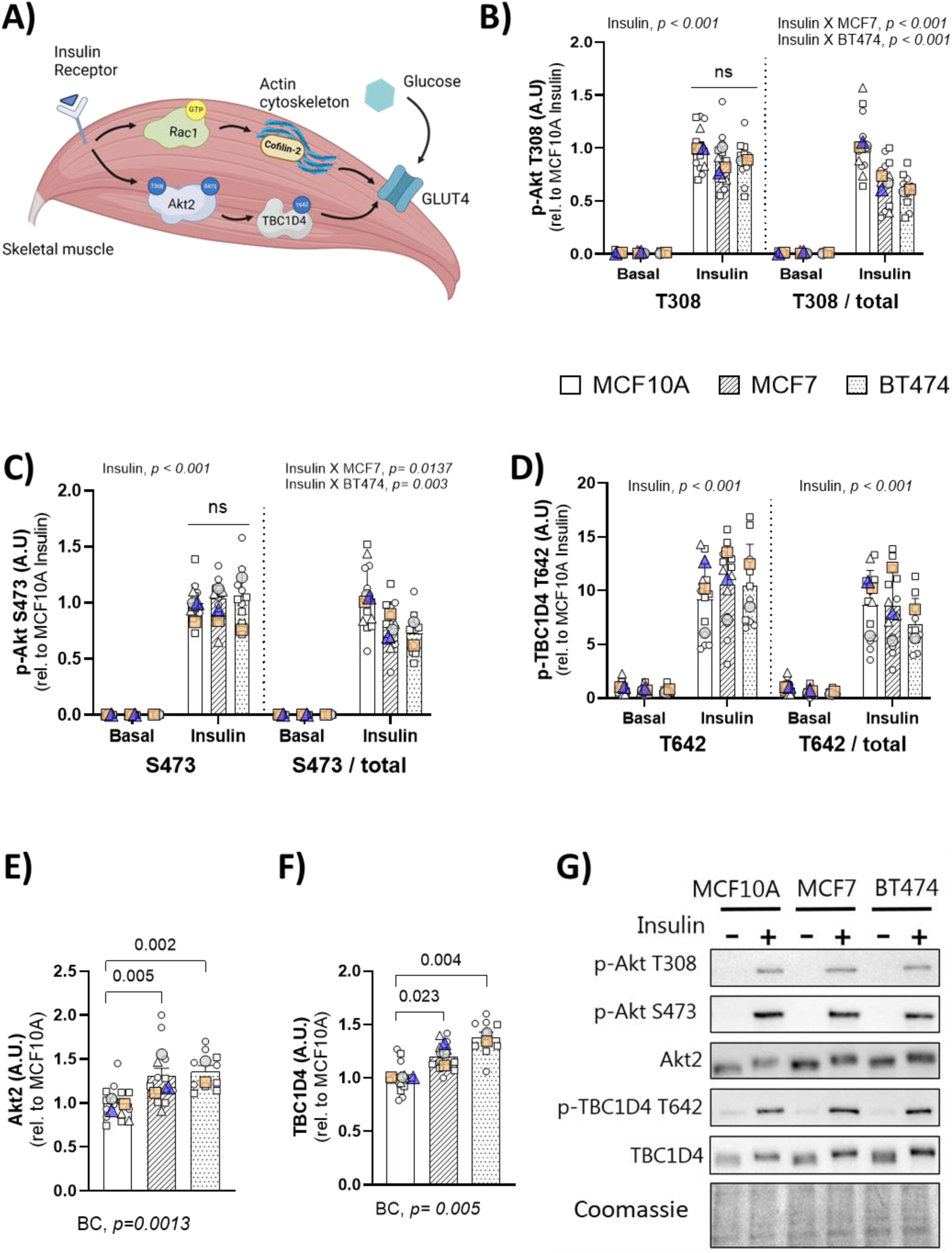
Breast cancer-secreted factors increase myotube Akt2 and TBC1D4 protein content without altering insulin-stimulated Akt-TBC1D4 phosphorylation. **A)** Schematic illustration of pathways analysed in GLUT4myc-L6 myotubes treated with breast cancer conditioned media downstream of the insulin receptor. Created with biorender.com. Quantifications of basal or insulin (10 nM, 25 minutes)-stimulated **B)** phosphorylated(p)-Akt T308, **C)** p-Akt S473, **D)** p-TBC1D4 T642, E) total Akt2, and **F)** total TBC1D4 protein level in GLLUT4myc L6 myotubes incubated with 40% conditioned media from MCF7 (striped), BT474 (dotted) and MCF10A (plain) cells for 72 hrs. N=3-5 repeated 2-3 times **G)** Representative blots including coomassie staining as loading control. Data are presented in super-plots as mean ±SEM incl. individual values that are colour- and symbol-coded (representing wells) matching each round (large color-/and shape-coded symbols representing the average of each experimental round)^44^. All data in (B, C, D) were evaluated with two-way ANOVA and in (E, F) were evaluated with one-way ANOVA using a pool of individual values from all three rounds. Main effects are indicated in the panels. Significant main effects and interaction (insulin X BC) in the ANOVAs were evaluated by Sidak’s post hoc tests: The exact p-values of breast cancer-conditioned media effect are stated on the graph when above p>0.001 or *** if below p<0.001. BC, breast cancer, A.U., arbitrary units, rel, related.

### Insulin-stimulated Rac1 activation and downstream signalling is dysregulated in muscle cells treated with breast cancer-conditioned media

With no detectable decrease in Akt-TBC1D4 signalling, we next turned to measure another critical signalling pathway regulating GLUT4 translocation, the small Rho GTPase Rac1^22–26,88^. Insulin-stimulated Rac1 activity (Rac1-GTP) was markedly reduced by conditioned media from MCF7 (−40 %, *p=0.0035*) and BT474 (−35%, *p=0.0033*) compared to control (Fig 5A). Rac1 activity at baseline was statistically indifferent between the groups, despite the numeric decrease of −25% (*p>0.10)*. The total protein level of Rac1 and its upstream inhibitor RhoGDIα^27^ were not affected by MCF7 or BT474 conditioned media (Fig 5B, C).

**Figure 5.**
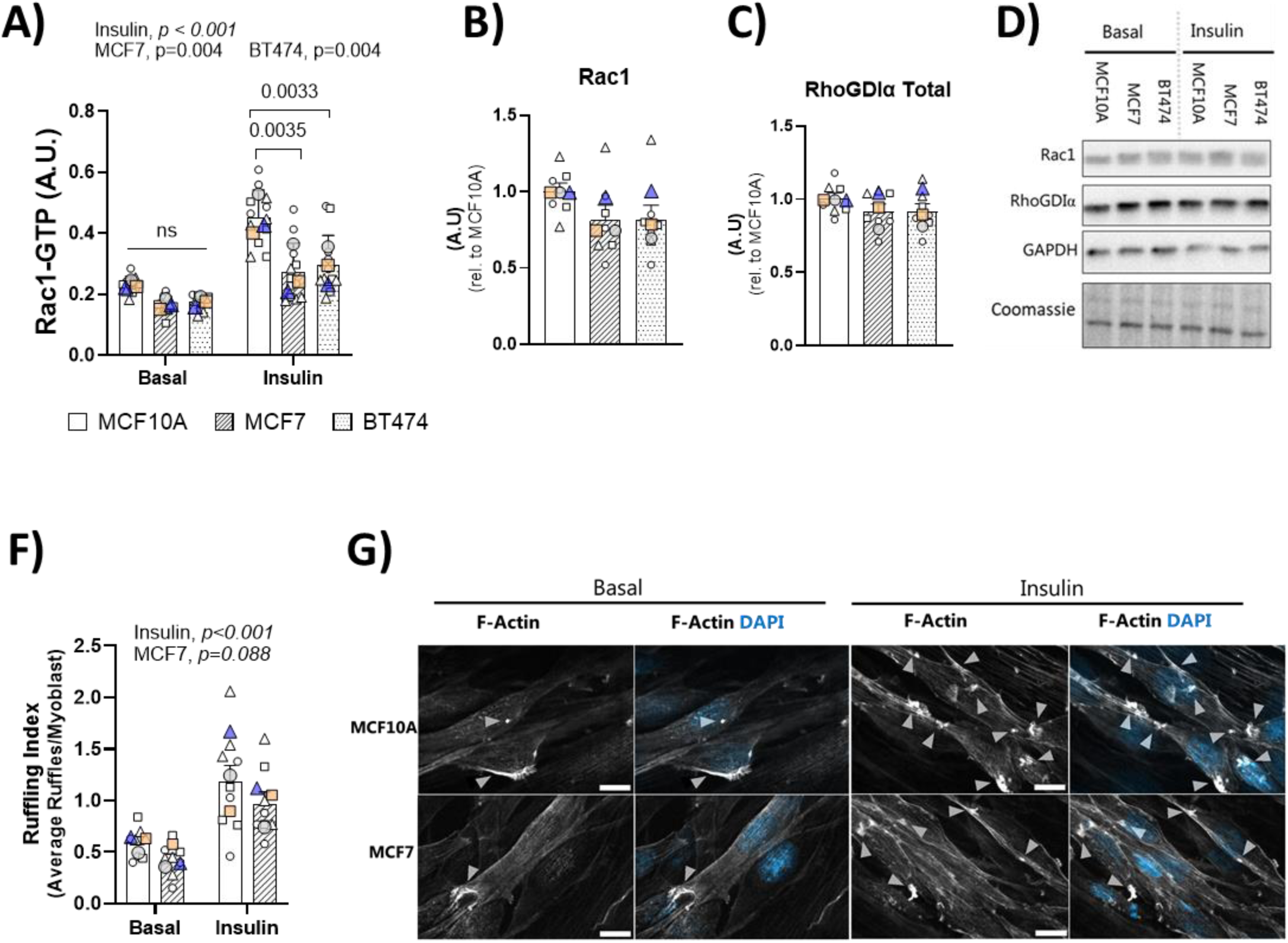
Insulin-stimulated Rac1 activation and downstream signalling is dysregulated in muscle cells treated with breast cancer-conditioned media. **A)** Basal and insulin-(10 nM, 10 min)-stimulated GTP bound Rac1 activity measured as GTP-bound Rac1 (Rac1-GTP) in GLUT4myc-L6 myotubes treated with 40% MCF7 (striped), BT474 (dotted) and MCF10A (plain) conditioned media for 72 hr. N=4 repeated three times. Quantification of total **B)** Rac1, **C)** RhoGDIα, and **D)** Cofilin-2 protein level in GLUT4myc-L6 myotubes incubated with 40% conditioned media from MCF7 (striped), BT474 (dotted) and MCF10A (plain). **E)** Quantifications of basal or insulin (10 nM, 10 minutes)-stimulation of phosphorylated (p)- Cofilin S3 in GLUT4myc-L6 myotubes incubated with 40% conditioned media from MCF7 (striped), BT474 (dotted) and MCF10A (plain) cells for 72 hrs. N=2-3 repeated three times. **F)** Representative blots including control protein staining by coomassie and GAPDH as housekeeping gene. **G)** Membrane ruffling index and **H)** representative immunofluorescence images of GLUT4myc-L6 myoblasts incubated with 40% conditioned media from MCF7 (striped), and MCF10A (plain) during basal and insulin (10 nM, 10 minutes)-stimulation using co-staining for F-Actin (Gray-scale), and DAPI (blue; nuclear stain). Scalebar = 100 µm at 63X magnification. N=3 repeated thrice. Data are presented in super-plots as mean ±SEM incl. individual values that are colour- and symbol-coded (representing wells) matching each round (large color-/and shape-coded symbols representing the average of each experimental round)^44^. All data in (A,D) were evaluated with two-way ANOVA and in (B,C,E) were evaluated with one-way ANOVA using the mean values of each round. Main effects are indicated in the panels. Significant main effects and interaction in the ANOVAs were evaluated by Sidak’s post hoc tests: The exact p-values of breast cancer-conditioned media effect are stated on the graph when above p>0.001 or *** if below p<0.001. ns; not significant, A.U., arbitrary units, rel, related.

Insulin-stimulated activation of Rac1 is required for insulin-induced cortical actin ruffling formation^89^, which is crucial for the fusion of GLUT4 vesicles with the plasma membrane^23^. Therefore, we measured the ruffling index (average number of ruffles per cell) with and without insulin stimulation (Fig. 5F, G). Given the similar mechanisms between MCF7 and BT474, we focused our ongoing efforts on MCF7 due to its more pronounced effect on insulin-induced GLUT4 translocation. In addition, cortical actin remodelling is more prevalent in myoblasts than in myotubes^90^. Therefore, we measured membrane ruffle formation in myoblasts to better understand the implications of breast cancer-conditioned media on skeletal muscle actin cytoskeleton remodelling. Insulin stimulation increased the number of ruffles per myoblast (MCF10A: 1.27±0.40, and MCF7: 0.96±0.34) within both conditions (*p<0.001*). We observed that the ruffling index tended (*p=0.088*) to be reduced by −25% in myoblasts treated with MCF7 conditioned media during both basal and insulin stimulation compared to MCF10A conditioned media. These results align with our findings from the proteomic analysis showing deregulated proteins related to actin cytoskeletal remodelling and Rho GTPase signalling (Fig. 1B-D). Altogether, our results suggest a potential role of Rac1-mediated signalling as a defective mechanism with breast cancer that could underlie the reduced GLUT4 translocation and glucose uptake.

Collectively, these results suggest that breast cancer-secreted factors induce changes in the proteomics and activates inflammation through activation of the TNFα-NF-ĸB-IL6 pathway, which further impacts sarcomere assembly, contractile function and myogenic differentiation. Ultimately, this leads to disorganisation of the cortical actin cytoskeleton via inhibition of the Rho GTPase Rac1 signalling, thus affecting the GLUT4 translocation and glucose uptake in skeletal muscle myotubes (Fig. 6).

**Figure 6.**
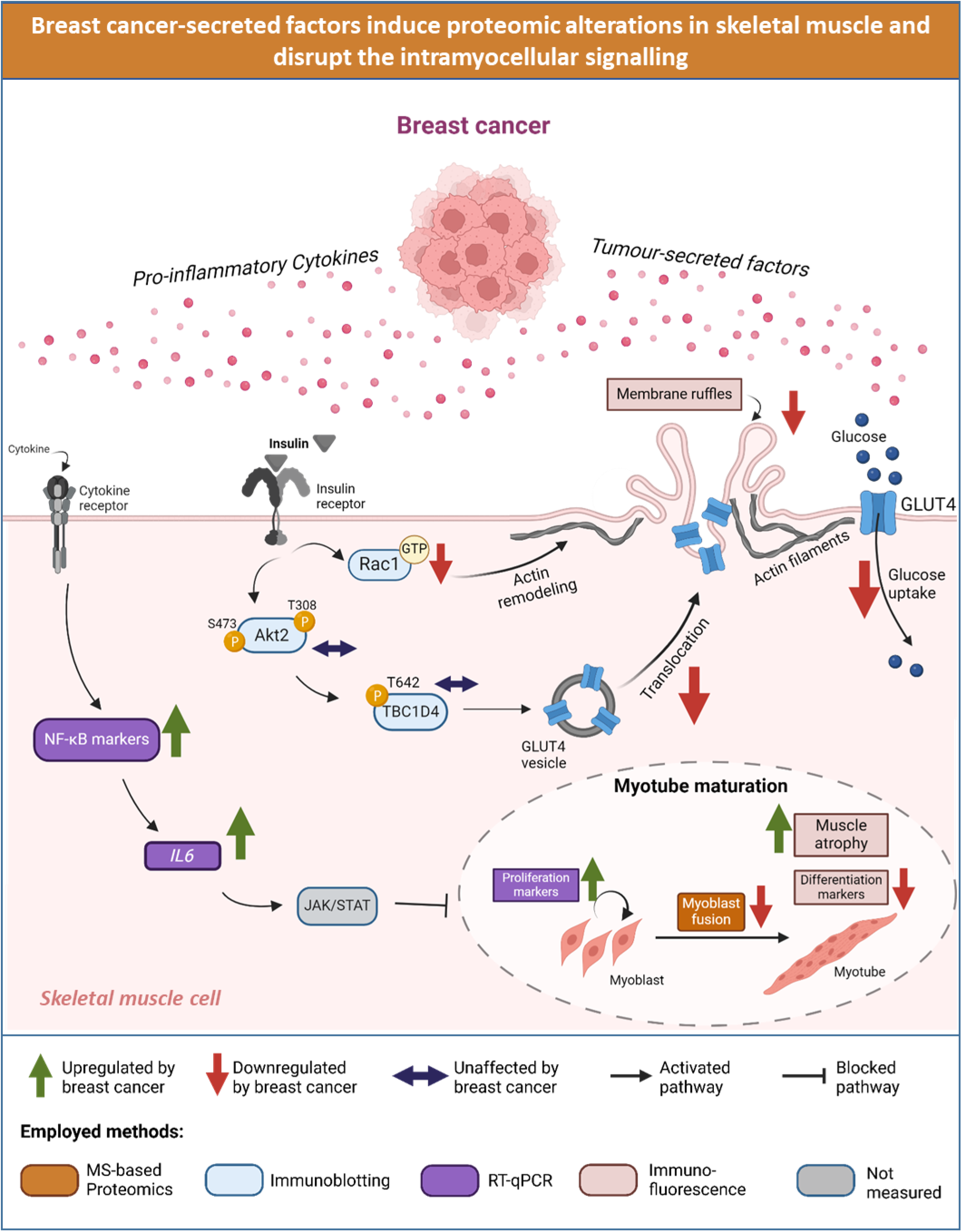
Schematic illustration of findings obtained in current study. Breast cancer-secreted factors induced proteomic alterations in pathways involved in sarcomere organization, myogenic programming, actin cytoskeletal remodelling, and vesicle trafficking. It also activated an inflammatory response via NF-κB pathway, and promoted muscle atrophy. Concomitantly, basal and insulin-stimulated GLUT4 translocation and glucose uptake were decreased. Insulin signalling via Rac1 signalling was blocked, whereas Akt-TBC1D4 pathway was conserved. JAK/STAT; Janus kinases/signal transducers and activators of the transcription. MS-based proteomics; mass spectrometry-based proteomics. RT-qPCR; Real-Time Quantitative Polymerase Chain Reaction. Illustration created by biorender.com.

## Discussion

The current study demonstrates that factors secreted from breast cancer cells directly influence skeletal muscle intracellular mechanisms. We observed altered muscle proteome, impaired myogenesis regulation, inflammation, and a lowering of basal and insulin-stimulated GLUT4 translocation and glucose uptake, likely driven by a reduction in insulin-activated Rac1-signalling. To our knowledge, these findings are the first to suggest that breast cancer-secreted factors may directly rewire skeletal muscle global proteome, myogenesis, glucose uptake, and insulin signalling, potentially explaining the high risk of metabolic dysfunctions in patients with breast cancer. Four key findings support this paradigm. First, myotubes exposed to breast cancer cell-conditioned media displayed marked proteomic changes. Second, breast cancer cell-conditioned media impaired myogenic differentiation, and triggered an inflammatory response through activation of the NF-κB pathway, inducing muscle atrophy. Third, breast cancer cell-conditioned media reduced basal and insulin-stimulated GLUT4 translocation concomitant with lower myotube glucose uptake. Finally, while canonical insulin signalling via Akt-TBC1D4 was unaffected by breast cancer cell-conditioned media, insulin-induced activation of Rac1 was blocked, and Rac1 signalling was disrupted.

Our first key finding was that breast cancer cell-conditioned media led to marked proteomic alterations in the myotubes. A recent study also revealed changes in the global proteome of primary human muscle cells incubated with breast cancer cell-conditioned media^34^. However, that study investigated the effect of the most aggressive triple-negative breast cancer subtype on muscle cells derived from a patient with type 2 diabetes. Similarly, more recent findings indicate significant changes in transcriptomic, metabolomics, and lipidomics in the skeletal muscle of immunodeficient breast cancer mice with triple-negative subtype^32^. To our knowledge, the effect of the hormone receptor-positive luminal A and luminal B breast cancer subtypes on skeletal muscle proteomics has not yet been investigated. This is important, as patients with hormone receptor-positive subtypes account for 70% of breast cancers, have the highest survival rates and experience a high prevalence of metabolic dysfunctions^5,6,91^. Although cancer treatment *per se* can induce metabolic alterations in patients with breast cancer, evidence shows that skeletal muscle from patients with breast cancer exhibits metabolic reprogramming independent of treatment^92^. This suggests that cancer-secreted factors directly affect skeletal muscle via alternative mechanisms to treatment.

Our global proteomic analysis revealed alterations in the myogenic programming and inflammation. We observed altered expression of proteins involved in cell cycle exit, CSRP3^93,94^ and PDLIM5^50,95^, myoblast fusion, SERPINE2^51^, and cytokine-mediated pro-inflammatory pathway, JAK2, TNFSF18, and COX2. Furthermore, pathways related to the contractile machinery, sarcomere organization, and actin filament binding were altered. Importantly, proteins involved in Rho GTPase signalling, IRSp53^52^ and STARS^53^, were downregulated. Rho GTPases, including Rac1, Cdc42 and RhoA, promote important cellular functions such as cell cycle progression, vesicle trafficking, glucose uptake, and actin cytoskeletal organization^96–98^.

In line with the proteomics results, our second key finding was that breast cancer-secreted factors inhibited myogenic differentiation, activated an inflammatory response via NF-κB and IL6, and induced muscle atrophy. Unlike the triple-negative breast cancer subtype, which was found to increase myogenic differentiation^33^, we observed that conditioned media from hormone receptor-positive subtypes inhibited myogenic differentiation. This is evident by our observation of reduced MHC expression, measured by immunofluorescence, and increased gene expression of the proliferation marker, *Mki67*. In consistency with the observations from the proteomic analysis, these findings suggest that breast cancer-secreted factors inhibit cell cycle exit, and block myoblast fusion.

In addition, breast cancer-secreted factors enhanced the inflammatory response via the NF-κB and Type 1 IFN pathways. The TNFα-mediated NF-κB pathway induces muscle atrophy through activation of IL6 and Janus kinases/signal transducers and activators of the transcription (JAK/STAT) pathway ^74,99^. We observed enhanced *Il6* gene expression, and an upregulated JAK2 protein abundance by proteomics analysis, which indicates a possible activation of the JAK/STAT pathway. Accordingly, myotubes exposed to breast cancer-secreted factors displayed reduced myotube width and concomitant MHC fragmentation, both considered markers for skeletal muscle atrophy ^79^. The inflammatory-induced skeletal muscle atrophy and cancer-related cachexia are both regulated by JAK/STAT pathway through the activation of STAT3 ^62,63,100,101^. Although cancer cachexia is rarely associated with breast cancer, several studies have demonstrated induced muscle weakness and inflammation by breast cancer in cells, mice, and humans^29,31–34,69,102–104^. Altogether, our results suggest that disrupted contractile machinery via sarcomere disorganization, and increased inflammatory-induced muscle atrophy could be the underlying causes of muscle weakness experienced with breast cancer.

Our third major finding that breast cancer cell-conditioned media lowered GLUT4 translocation and glucose uptake in GLUT4myc-L6 myotubes could be linked to the increased NF-κB pathway observed in myotubes treated with breast cancer conditioned media. Notably, enhanced NF-κB activation reduces glucose uptake and GLUT4 translocation in C2C12 myotubes^77^. We speculate that the direct effects of factors secreted by breast cancer cells could contribute to the metabolic perturbations observed in up to 60% of patients with breast cancer^5–8^. This is supported by a growing number of studies reporting that breast cancer patients experience significant metabolic dysregulations, including hyperglycaemia, and hyperinsulinemia^6–8,38^. In fact, patients with cancer, including breast and lung cancer, display an increased risk of developing diabetes after their diagnosis^9,10^. Additionally, a study from our group has shown that Lewis lung cancer caused metabolic perturbations affecting glucose tolerance and skeletal muscle insulin-stimulated glucose uptake in mice^35–37^. Such studies suggest that metabolic regulation is disturbed in several cancers.

Our final major finding was that breast cancer cell-conditioned media rewired the intramyocellular insulin signalling by selectively blocking Rac1 activity. Both MCF7 and BT474 conditioned media did not affect the key insulin-responsive pathway led by Akt, and its downstream protein TBC1D4. Although one study showed a reduced basal p-Akt S473 in primary human myotubes exposed to MCF7 conditioned media^34^, the functional impact on insulin signalling was not investigated. Notably, we observed an upregulated total Akt2 and TBC1D4 protein content with breast cancer cell-conditioned media, suggesting a compensatory response to rescue the effect of breast cancer-conditioned media. These results align with previously reported *in vivo* studies showing an increased total Akt2 and TBC1D4 protein content in Lewis lung cancer tumour-bearing mice^35,36^.

While Akt-signalling was conserved, breast cancer-secreted factors markedly reduced GTP-bound Rac1. The Rho GTPase Rac1 plays a major role in the regulation of skeletal muscle insulin sensitivity and glucose uptake^22–26,88^, and induces GLUT4 translocation by promoting actin cytoskeletal remodelling in response to insulin^105^. More specifically, Rac1 induces cortical actin cytoskeletal remodelling by promoting the formation of membrane ruffling at the plasma membrane^80^. Disrupted cortical actin cytoskeletal remodelling prevents the insertion of GLUT4 into the plasma membrane^80^. Our proteomic results indicated a marked alteration in proteins involved in actin cytoskeleton organization. We observed reduced membrane ruffling with MCF7 conditioned media, aligning with the decreased Rac1 activity. Rac1 is implicated in several pathological conditions, making it a strong candidate for therapeutic target ^25,27,106–108^. Besides insulin signalling, Rac1 also regulates key proteins involved in skeletal muscle myogenesis^109,110^, suggesting a potential role of Rac1 in our current observations of inhibited myogenic differentiation. Altogether, our results suggest that breast cancer-associated metabolic dysfunctions could, in part, be induced by abrogated skeletal muscle insulin signalling, likely via Rac1 signalling^111,112^.

Several studies agree on the association between breast cancer and metabolic dysfunction, but several confounding factors, such as ethnicity, physical activity, menopausal state, dietary intake and treatment, could play a role in this association^113^. Therefore, our *in vitro* approach investigating the link between breast cancer and skeletal muscle insulin sensitivity is independent of any confounding factors and brings novelty to the field to help elucidate the direct effect of breast cancer on skeletal muscle. However, we did not identify the factor(s) secreted by breast cancer cells that could induce insulin resistance in muscle, as our main objective was to address the intramyocellular changes caused by breast cancer-secreted factors. The secretome composition of conditioned media includes proteins, cytokines, growth factors, chemokines, lipids, and nucleic acids with various functions involved in cell-to-cell communication^114^. Notably, conditioned media from MCF7 breast cancer cells revealed four specific biomarkers, including IGFBPs, which are also identified in plasma from patients with breast cancer^46^.

Our present *in vitro* findings align with the documented results from mice and humans showing significant molecular rewiring of skeletal muscle with breast cancer^29–34,69^. Moreover, our study provides mechanistic evidence that might explain why women who develop breast cancer have a higher incidence of metabolic dysfunctions, such as diabetes, compared to women who do not develop breast cancer^92,115^. As up to 60% of women with breast cancer develop metabolic disorders during or following their treatment^8,10,26^, enhancing muscle health could be a powerful means to improve metabolic regulation for these women. Exercise training is the most effective means to improve skeletal muscle insulin sensitivity and strength^116^. Accordingly, exercise can prevent metabolic disturbances in breast cancer survivors^6,31,117,118^, and reduce the risk of breast cancer and breast cancer recurrence^119–122^. Furthermore, Rac1 activation is also upregulated by exercise ^123,124^, making it a potential target for treatment against breast cancer-induced metabolic dysfunctions.

## Conclusion

In conclusion, our study demonstrates that secreted factors from hormone receptor-positive breast cancer cells, MCF7 and BT474, alter muscle proteome in pathways related to myogenesis, inflammation, actin cytoskeleton organization, and vesicle trafficking, complementing the disrupted GLUT4 translocation, glucose uptake, and intramyocellular signalling as well as induced muscle atrophy. This disruption occurs through selective inhibition of Rac1 activation and activation of pro-inflammatory response via NF-κB pathway while preserving Akt signalling. These findings provide compelling evidence that factors secreted by breast cancer cells can directly alter skeletal muscle glucose uptake. This rewiring of skeletal muscle basal and insulin signalling underscores a potential mechanistic link between breast cancer and metabolic dysfunction, highlighting the need for further investigation into targeted therapies that address this metabolic interplay.

## Contributions

M.S.A. and L.S. conceptualized and designed the study. M.S.A. conducted the experiments, performed the laboratory analysis, and analysed the data. M.S.A. and L.S. wrote the manuscript. S.B.O., E.F., E.B., M.P., S.S.B.T., S.H.R., and S.D.J. all took part in conducting the experiments, performing laboratory analysis, and/or interpreting the data. All authors commented on and approved the final version of the manuscript. All authors are guarantors of this work and take responsibility for the integrity of the data and the accuracy of the data analysis.

## Acknowledgments

We thank Professor Amira Klip (Cell Biology Program, Research Institute, The Hospital for Sick Children, Toronto, Canada) for the kind donation of the GLUT4myc-L6 muscle cells. Mass spectrometry-based proteomics analyses were performed by the Proteomics Research Infrastructure (PRI) at the University of Copenhagen (UCPH), supported by the Novo Nordisk Foundation (NNF) (grant agreement number NNF19SA0059305). L.S. was supported by the Novo Nordisk Foundation (NNF16OC0023418 and NNF18OC0032082) and Independent Research Fund Denmark (9039-00170B). M.J. was supported by the Danish National Research Foundation (DNRF125 to M.J.). M.S.A. was supported by a research grant from the Danish Diabetes Academy, which is funded by the Novo Nordisk Foundation, grant number NNF17SA0031406.

